# Temperature-driven selection on metabolic traits increases the strength of an algal-grazer interaction in naturally warmed streams

**DOI:** 10.1101/104513

**Authors:** C. -Elisa Schaum, Bio244 Students, Richard ffrench-Constant, Chris Lowe, Jón S Ólafsson, Daniel Padfield, Gabriel Yvon-Durocher

## Abstract

Trophic interactions are important determinants of the structure and functioning of ecosystems. As the metabolism and consumption rates of ectotherms increase sharply with temperature, there are major concerns that global warming will increase the strength of trophic interactions, destabilizing food webs, and altering ecosystem structure and function. We used geothermally warmed streams that span a ∼10°C temperature gradient to investigate the interplay between temperature-driven selection on traits related to metabolism and resource acquisition, and the interaction strength between the keystone gastropod grazer, the wandering snail *Radix balthica*, and a common algal resource. Populations from a warm stream (∼28°C) had higher maximal metabolic rates and optimal temperatures than their counterparts from a cold stream (∼17°C). We found that metabolic rates of the population originating from a warmer stream were higher across all measurement temperatures. A reciprocal transplant experiment demonstrated that the interaction strengths between the grazer and its algal resource were highest for both populations when transplanted into the warm stream. In line with the thermal dependence of respiration, interaction strengths of grazers from the warm stream were always higher than those of grazers from the cold stream. These findings suggest that warming can increase the strength of algal-grazer interactions through the thermodynamic effects of higher temperatures on physiological rates as well as through correlated increases in *per capita* metabolism and consumption.

## INTRODUCTION

The strength of consumer-resource interactions (e.g. the effect of a consumer on the population density of its prey) plays a critical role in shaping the stability of food webs (May, 1973; Paine, 1980; McCann *et al.*, 1998; Otto *et al.*, 2007). Grazing is an important class of consumer-resource interaction, determining the flux of energy and materials from autotrophs to heterotrophs. There are currently major concerns that global warming will increase the impact of grazers on algal or plant communities because the ingestion and respiration rates of heterotrophs tend to be more sensitive to rising temperatures than rates of photosynthesis and growth in autotrophs (O’Connor, 2009; Gilbert *et al.*, 2014; West & Post, 2016). Stronger interactions have the potential to destabilise food webs and consequently, warming induced increases in interaction strengths could have fundamental implications for ecosystem structure and function. For example, elevated grazing rates in aquatic ecosystems, driven by the mismatch in thermal sensitivity between autotrophs and heterotrophs, are a key driver of projected declines in aquatic primary production over the 21^st^ century in models of ocean biogeochemistry (Laufkötter *et al.*, 2015).

The effects of temperature on metabolic rates and traits associated with consumer-resource interactions (e.g. attack rates, handling times) often follow characteristic unimodal thermal response curves, in which rates increase exponentially to an optimum and decline rapidly thereafter (Dell *et al.*, 2011; Englund *et al.*, 2011; Rall *et al.*, 2012; Dell *et al.*, 2014; Gilbert *et al.*, 2014). Integrating thermal responses for metabolism and interaction-traits with dynamical models of consumer-resource interactions offers a promising framework for predicting food web responses to global warming (Vasseur & McCann, 2005; Shurin *et al.*, 2012; Binzer *et al.*, 2015). However, thermal response curves are often flexible, and can shift when organisms are exposed to novel thermal environments, both via phenotypic plasticity and adaptive evolution (Angilletta *et al.*, 2003; Kingsolver *et al.*, 2004; Deutsch *et al.*, 2008;Kingsolver & Huey, 2008). Consequently, plasticity and evolution have the potential to modulate the effects of rising temperatures on the strength of species interactions (Sentis *et al.*, 2015). For example, if metabolic rates are down-regulated after long-term exposure to higher temperatures (Addo-Bediako *et al.*, 2000), then compensatory metabolic responses to warming could mitigate predicted increases in consumer-resource interaction strength. How these long-term responses to warming affect rates of metabolism and in turn, the strength of consumer-resource interactions, are largely unknown, limiting our ability to predict how trophic interactions will change in response to warming in the long-term.

There is evidence from studies across naturally occurring thermal gradients over large spatial scales, that local thermal adaptation can play an important role in shaping the strength of species interactions (Barton, 2011; De Block *et al.*, 2012). While these studies provide important insights into how consumer-resource interactions are shaped by evolution across thermal gradients (Fukami & Wardle, 2005), their usefulness for understanding responses to rapid climate warming might be limited, because other factors, such as day length, light intensity and precipitation, tend to be confounded with temperature along such broad scale spatial gradients. Furthermore, the timescales over which local adaptation has occurred in such broad scale studies could be much longer than the rapid evolutionary change required to keep pace with climate warming (Loarie *et al.*, 2009; Hoffmann & Sgrò, 2011). Here, we investigate how temperature-driven selection on traits that determine the thermal responses of metabolism and resource acquisition affect the strength of a keystone grazing interaction (the gastropod *Radix balthica,* which grazes algal biofilms in streams) in naturally warmed Icelandic geothermal streams spanning a gradient of 11°C. Critically, temperature is the main abiotic factor that varies among streams in the catchment and is not correlated with pH, conductivity or inorganic nutrient concentrations (see Table 1). These streams are thought to have been subject to geothermal heating for at least the last century (OGorman *et al.*, 2012).

**Table 1.** Physical and chemical characteristics of the streams. Temperature data were collected over a 3-day period. All other parameters were collected on the first day of the day of the experiment. Temperature data are displayed as means ± 1SD. All other data were originally collected for correlation with temperature across the catchment area (all 11 streams), so that replication was on the level of stream identitys

| Parameter | Stream 5 | Stream 11 |
| --- | --- | --- |
| Average temperature (°C, 5 days) | 17.5 ± 4.5 | 28.3 ± 1.3 |
| pH | 7.63 | 7.17 |
| Conductivity | 273.6 | 235.7 |
| NO<sub>2</sub> | 0.22 | 0.24 |
| NO<sub>3</sub> | 0.57 | 0.29 |
| NH<sub>4</sub> | 0.17 | 0.19 |
| PO<sub>4</sub> | 0.27 | 0.35 |

This system therefore provides the opportunity to investigate how long-term differences in temperature between otherwise similar sites shape the expression of metabolic traits and the subsequent impact of any temperature-driven selection on species interactions in a natural system. Specifically, we ask: can long-term differences in temperature drive selection for metabolic traits that attenuate the direct effects of warming on the strength of consumer-resource interactions?

## METHODS

### Study site

The streams are located North of the Hveragerði valley, in the south east of the Hengil high temperature geothermal field, Iceland (N64° 0’ 2.944" W21° 11’ 17.451") and consist of a catchment of 11 streams spanning a temperature gradient of approximately 20 °C (see Figure 1 and SI Figure 1). Two streams, stream 5 (17.5 °C ± 4.5 °C, hereafter ‘cold stream’) and stream 11A (28.3 °C ± 1.3 °C, hereafter ‘warm stream’, see Table 1 for a comparison of other chemical and physical parameters), were chosen for experiments due to their close proximity to each other, the large temperature differential and similar abundances of the keystone grazer, *Radix balthica*. The grazer plays an important functional role in geothermal stream ecosystems, where grazer biomass as well as grazing rates are strongly influenced by temperature (OGorman *et al.*, 2012). The two streams are similar in all other measured physical and chemical characteristics but differ in average temperature by 11 °C (see Table 1), and hence present an opportunity to investigate how the effects long-term differences in temperature shape consumer-resource interactions.

**FIGURE 1.**
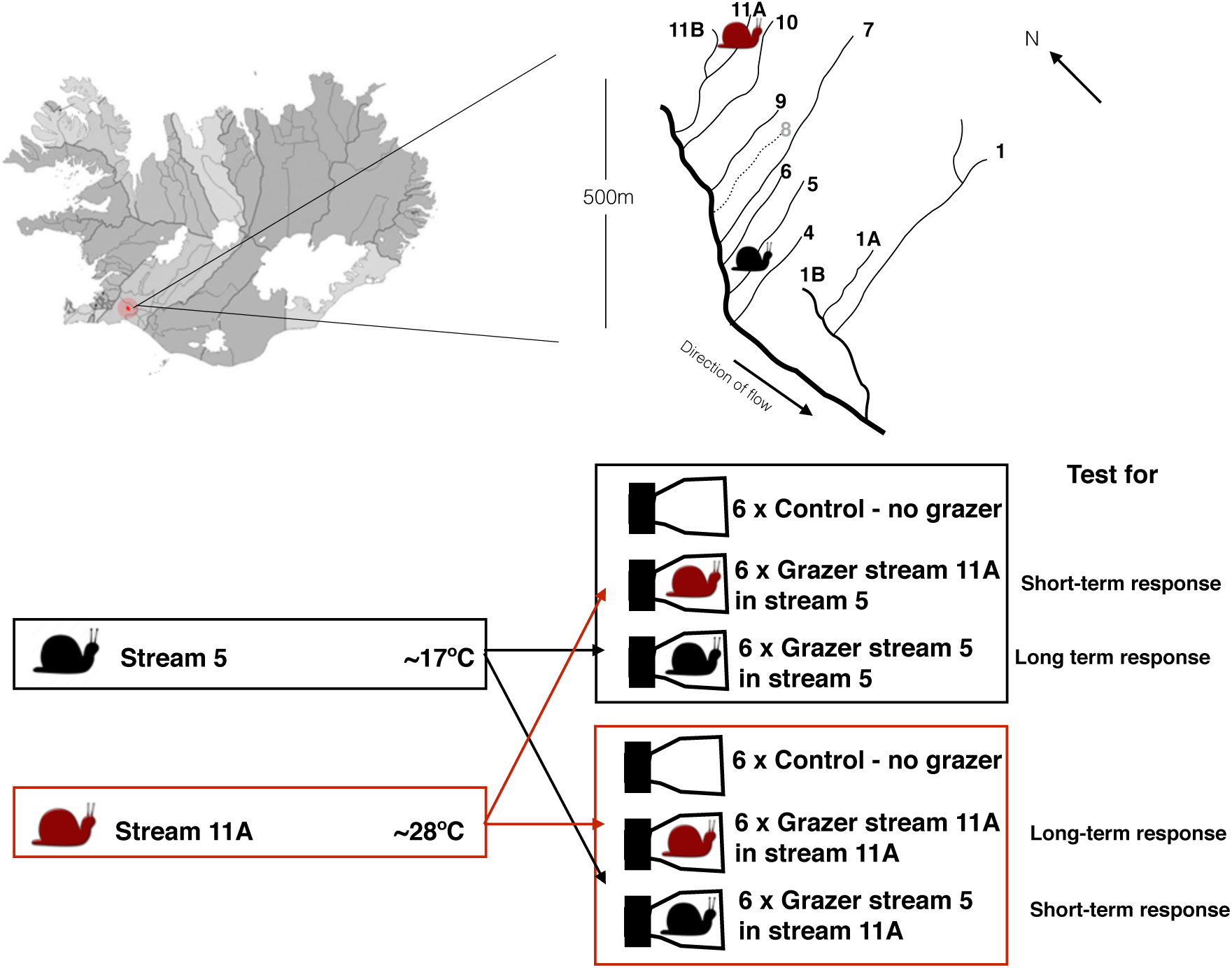
Map and experimental set-up. **Top panel**: The catchment area, with streams used in this experiment indicated by black (for the colder stream 5 with 17.5 °C ± 4.5 °C) and red (for the warmer stream 11A with 28.3 °C ± 1.3 °C) snail icons. **Lower panel**: Schematic overview of experimental set-up for the grazing experiment.

### Grazer metabolism

To quantify whether the different thermal regimes in the two adjacent streams resulted in divergence in metabolic traits of *R. balthica* we measured the acute responses of respiration to a broad gradient in temperature. We collected 33 individuals of similar weight and length from each stream, which were cleaned from any algal debris to avoid carry-over of a food source into the tank or subsequent respiratory measurements on the oxygen electrode. The snails were kept overnight in aerated tanks at the average stream temperature of origin and in the absence of a food source to minimise any potential effects of differences in food quantity or quality between streams. Respiration was quantified as the rate of oxygen consumption in a Clark-Type oxygen electrode, measured between 4 – 44 °C in 4 °C increments (11 temperatures in total). At each temperature, respiration was measured for 3 individuals, and a different set of individuals was measured at each temperature (i.e. each animal was only subjected to a single assay). Individuals were allowed 15 minutes at the assay temperature prior to the measurements. The subsequent thermal responses of respiration were quantified using a modification of the Sharpe-Schoolfield equation (see (Schoolfield *et al.*, 1981) and (Sharpe & DeMichele, 1977)for the original equation):

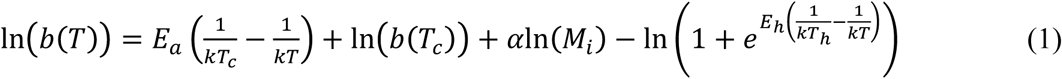

where *b*(*T*), is the *per capita* metabolic rate (µmol O_2_ L^-1^ h^-1^) at temperature *T* in Kelvin (K), *k* is Boltzmann’s constant (8.62×10^-5^ eV K^-1^), *E*_*a*_ is an apparent activation energy (in eV) for the metabolic process, ln (*b*(*T*_*c*_)) is the rate of metabolism normalised to an arbitrary reference temperature, *T*_*c*_ = 18 °C, where no low or high temperature inactivation is experienced. *M*_*i*_ is the mass (g) of an individual *i*, α is the allometric scaling exponent that characterises the power-law relation of mass and metabolic rate (Brown *et al.*, 2004). *E*_*h*_ characterizes temperature-induced inactivation of enzyme kinetics above *T*_*h*_ where half the enzymes are rendered non-functional. Differentiating equation (1) and solving for the global maxima yields an expression for the optimum temperature

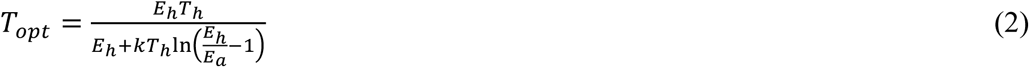

Equation (1) differs from the Sharpe-Schoolfield equation (Sharpe & DeMichele, 1977; Schoolfield *et al.*, 1981) in a number of ways. First, we account for the power law relation between body mass and metabolic rate, *M*^*α*^ (Brown *et al.*, 2004). Second, we exclude parameters from Eq. (1) used to characterize low-temperature inactivation due to insufficient data to quantify this phenomenon in our analysis. Third, rather than characterize temperature effects below *T*_opt_ using the Eyring (1935) relation, 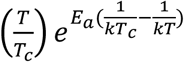, we instead use the simpler Boltzmann factor, 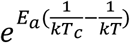 This simplification enables an explicit solution for *T*_opt_ (Eq. 2) and facilitates more direct comparison with previous work on the temperature dependence of metabolism using metabolic theory e.g. (Allen *et al.*,2005, Gillooly, 2001; Brown *et al.*, 2004; Van M Savage *et al.*, 2015).

The parameters, ln *b*(*T*_*c*_), α, *E*_*a*_, *E*_*h*_, *T*_*h*_, and *T*_opt_, in Eqs. (1) & (2) represent traits characterising the metabolic thermal response that we expect to be under selection in *R. balthica* inhabiting the hot and cold streams. We tested for differences in each of the parameters between the populations of *R. balthica* by fitting the respiration data to Eq. (1) using generalised non-linear least squares regression (within the ‘gnls’ function in the ‘nlme’ package for R, package version 3.1-128) and including ‘origin’ as a two level factor (i.e. ‘cold’ and ‘warm’ stream). We tested for differences between populations for each parameter by sequentially removing the effect of ‘origin’ on each parameter and comparing the Akaike information criterion for small sample sizes (AICc) for all possible models (see SI Table 1 and SI Table 2) using the ‘aictab’ and ‘modavg’ functions from the AICcmodavg package (package version 2.1-0). The model chosen for further exploration was that with the lowest (AICc) value. Model averaging was carried out when models fell within 2 AICc units of each other, and the conditional averages of the parameters were used for curve fitting and interpretation (see also Table 2). The relative importance of the fixed factors in the averaged model was determined using the sum of their relative weights.

**Table 2.**
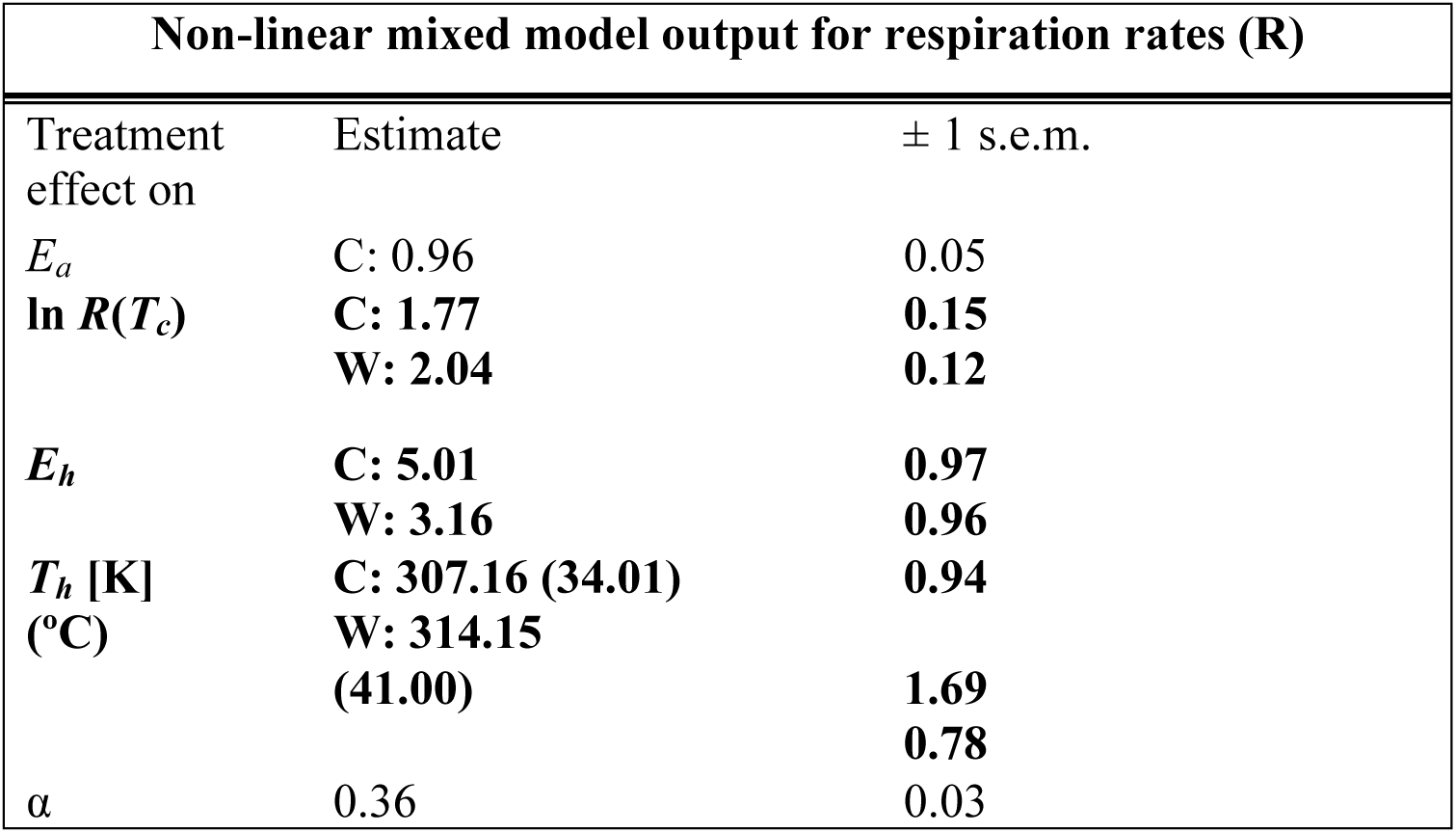
Parameter estimates and output from the best fitting gnls model to the thermal response curves of respiration rates. Differences in treatments are given in **bold.** Parameter estimates are taken from the averaged generalised linear models along with their standard errors (± 1 s.e.m). C = cold stream. W = warm stream. See Supporting Information for details on model selection and information on AICsc scores for all possible models. Here, the model average of the conditional average output for the four best models (within 2 AICc units of each other) is displayed.

### Reciprocal transplant experiment

We carried out a reciprocal transplant experiment to determine how long-term differences in temperature and the resultant impacts on metabolic traits affect the strength of algal-grazer interactions. We achieved this by placing snails from each population in microcosms consisting of a tissue culture flask on which diatom biofilms had been established. Diatoms of the genera, *Acnanthes, Nitzschia, Navicula*, and *Gomphonema* are common in streams across the Hengill volcanic area (Gudmundsdottir *et al.* 2013)and were ordered from culture collections (Culture collection of algae and protozoa and Sciento) and grown in the laboratory in mixed assemblages to yield common resource for testing the effects of temperature and local adaptation on grazing. The diatom assemblages were inoculated into Corning plastic translucent flasks (maximum volume 1L) with 20 mL COMBO medium (Kilham *et al.*, 1998), and brought to a salinity of 5-10 (equivalent to approximately 5-10 g salts/kg water) to match the slightly elevated salinity and conductivity found in these thermal stream environments (Gudmundsdottir et al. 2013). The flasks were turned onto their sides to allow for a larger area of biofilm growth on the base (∼ 60 cm^2^ in total per flask) and the algal communities were left to grow for 14 days prior to the experiment. After 14 days, all flasks had substantial biofilm development on the base and were used as microcosms for the *in situ* reciprocal transplant experiment. Analysis of control flasks (no grazer) showed that growth of the diatom lawn *per se* did not differ significantly for flasks placed in hot or cold streams (SI Figure 2, one-way ANOVA *F*_1,10_ = 1.28, *P* = 0.26). Thus, any changes to the biofilm biomass in the experiment can be attributed to the per capita effects of the grazer.

**FIGURE 2.**
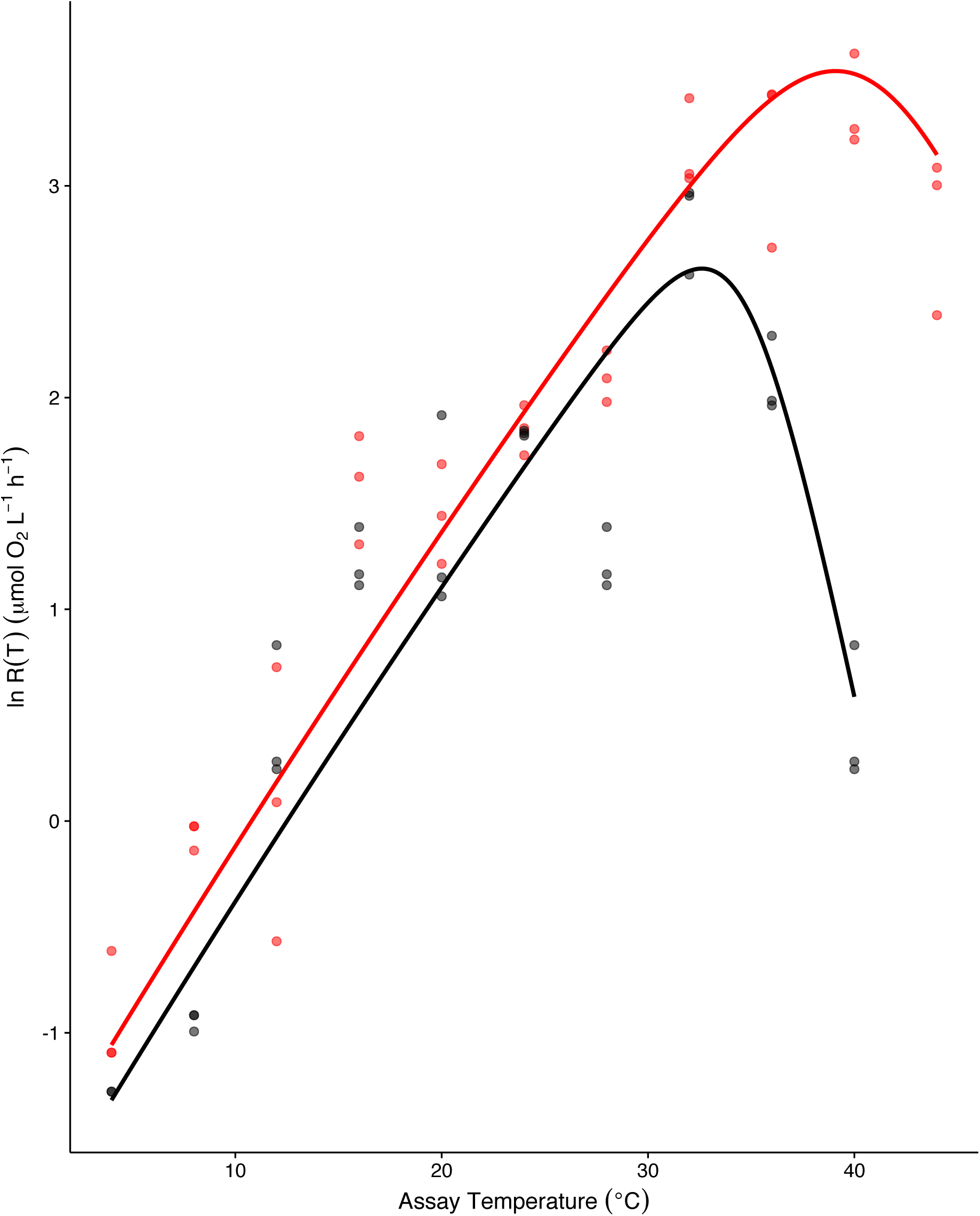
Thermal response curves for respiration. Thermal response curves of respiration rates in µmol O2 L^-1^ h^-1^ as a function of increasing temperature for populations of the snail *Radix balthica* from the cold (black) and warm (red) stream. Lines are derived from fitting a modified Sharpe-Schoolfied equation (see methods) to the rate data. Snails from the warm stream have higher temperature normalised metabolic rates (ln *R*(*T*_*c*_), with T_c_=18°C) at all measurement temperatures and have higher optimal temperatures (*T*_opt_), than snails from the cold stream. The inactivation energy (*E*_*h*_) is lower in snails from the warm stream, resulting in a curve that is both broader and elevated in comparison to the thermal response curve of respiration for snails from the cold stream.

The experiment consisted of 3 treatments (each with 6 replicate microcosms placed in each of the 2 streams): (i) a control microcosm in which a biofilm was present and no *R. balthica* were added, (ii) an ‘origin’ treatment in which *R. balthica* that were resident in the stream were added to microcosms, and (iii) a ‘transplanted’ treatment in which *R. balthica* that were from the adjacent stream were added to microcosms. *R. balthica* individuals were collected from the 2 streams the day before the experiment and were starved for 24h in the laboratory in aerated tanks at the average temperature of the stream of origin. There was no significant difference in average snail weight between the two streams (see SI Figure 3, one-way ANOVA: *F*_1,408_ = 0.15, *P* = 0.7). Microcosms were assembled by adding 3 snails of similar body dimensions (0.35 ± 0.03 g of *R. balthica* weight reported as blotted fresh weight throughout) and 100 mL of 0.4 µm filtered water from the stream in which the microcosm was to be placed. This resulted in a grazer density of 5 individuals m^-2^, which was comparable to the average *in situ* density in the streams (see SI Figure 4, no significant difference in in situ density between the two-streams: one-way ANOVA: F_1,66_, P = 0.54).This design was preferred to a set-up with each microcosm holding a single grazer, which attempt to exclude the effects of mutual interference on feeding behaviour e.g. (Skalski & Gilliam, 2001; Rall *et al.*, 2009; Lang *et al.*, 2011; Vucic-Pestic *et al.*, 2011), because (i) the experimental densities are representative of natural conditions; and (ii) the consumption rates of a single individual were insufficient to detect a significant change in algal biomass. The microcosms were submerged in each stream and the snails were left to graze for 48 hours.We observed no grazer mortality over the experimental period.

### Interaction strength

At the end of the experiment, algal biomass in each of the microcosms was quantified via methanol chlorophyll extraction modified from (Holm-Hansen & Riemann, 1978). Here, the walls of the microcosms were scrubbed until all biofilm particles were in suspension. The solution was filtered onto a 0.4µm GF/F filter, which was then ground in methanol for 5 minutes. The samples were centrifuged at 3500 rpm for 15 minutes and the absorbance of the supernatant was measured at 632nm, 665nm, and 750nm. Total chlorophyll content in µg mL^-1^ was then calculated as described in Holm-Hansen & Riemann (1978). The *per capita* interaction strength in each microcosm was then estimated by calculating the dynamic index (DI, see also (Berlow *et al.*, 2004)for a technically similar set-up):

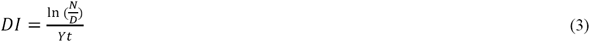

where DI is the dynamic index (g C^-1^ h^-1^), *N* is total chlorophyll (sum of Chl *a* + Chl *c*) content of control, *D* total chlorophyll in the grazed microcosm, *Y* is the grazer biomass (g C), and *t* is time in hours. Snail blotted wet weight was converted to carbon mass (in grams) using conversion factors that assume dry weight to be 7.5% of the blotted wet weight (Ricciardi & Bourget, 1998) and a carbon content of 22% dry weight (Burgmer *et al.*, 2010).

We carried out two analyses using the data from the reciprocal transplant experiment. The first analysis, used a generalised linear model (GLM), with ‘interaction strength’ as the response variable and ‘origin’ (‘cold’ or ‘warm’ stream) and ‘transplant temperature’ (17.5 and 28.3 °C) as potentially interacting factors. We used this analysis to determine (i) whether interaction strengths differed between snails that originated from the warm or cold streams (e.g. a main effect of ‘origin’); (ii) whether interaction strengths were temperature dependent (e.g. a main effect of ‘temperature’); and (iii) whether the temperature dependence of interaction strength differed between the snails from the cold and warm streams (e.g. interaction between ‘origin’ and ‘temperature’).

The design of the reciprocal transplant experiment also enabled us to disentangle short-term temperature responses attributable to acclimation (e.g. responses to the temperature in the ‘transplanted’ stream) from those reflecting processes operating over longer, time scales (e.g. adaptation to the stream of ‘origin’). Note that these ‘long-term’ effects, which we call ‘adaptation’, could reflect strict genetic microevolution (e.g. resulting in divergent genotypes among populations) or they could represent non-genetic effects of the different temperature regimes that manifest over ontogenetic development, but are nevertheless adaptive (Bonduriansky *et al.*, 2011). In the second GLM we included ‘interaction strength’ as the response variable and ‘timescale’ (‘short’ or ‘long’) and ‘transplant temperature’ (17.5 and 28.3 °C) as potentially interacting factors. Here, ‘short-term’ temperature responses were characterised as the change in interaction strength between the stream of origin and the transplant stream. By contrast, the ‘long-term’ temperature response was characterised as the change in interaction strength comparing measurements made only when the snails were in their stream of origin. For better comparison of the steepness of the respiration reaction norms, we re-express the transplant temperature data as Boltzmann temperatures 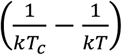 so that the coefficients of the model yield activation energies in units of eV (see Eq. (1)). In this analysis, a significant interaction between ‘transplant temperature’ and ‘timescale’ would demonstrate that the temperature dependence of interaction strength differs between the ‘short-term’ (*E*_short_, change in interaction strength between the stream of origin and the transplant stream, see also Fig. 3), and ‘long-term’ (*E*_long_, i.e. change in interaction strength comparing measurements made only when the snails were in their stream of origin, see also Fig. 3). We assume that *E*_short_ captures rapid physiological plasticity (e.g. acclimation) in interaction strength in response to a change in temperature and *E*_long_ captures processes operating over longer timescales – e.g. genetic microevolution and non-genetic developmental effects. Consequently, the component of the temperature sensitivity attributable to ‘adaptation’ (recognising that this might be genetically and/or developmentally determined) is given by *E*_adapt_ = *E*_long_ - *E*_short_.

**FIGURE 3.**
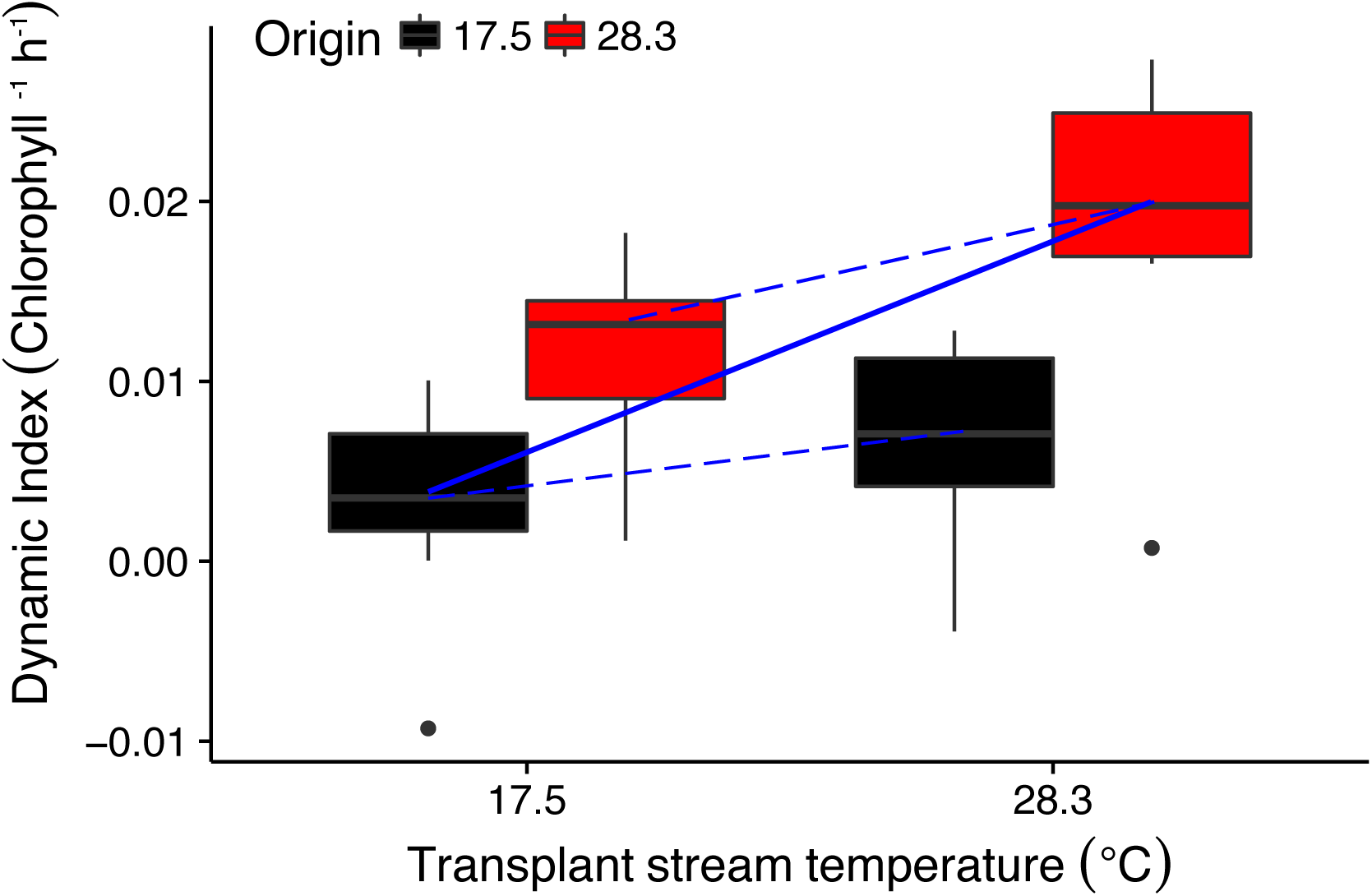
Long-term and short-term effects of stream temperature on interaction strength. Long-term and short-term effects of temperature in interaction strength measured via the dynamic index in units of chlorophyll consumed per hour. Populations originating from the warm stream have stronger interaction strength indices in all environments and the highest dynamic index overall was found for snails from the warm stream in their original environment. Interaction strength increased with temperature both in the short-term (*E*_short_, dashed blue lines) and in the long-term (*E*_long_, solid blue line), with *E*_long_ significantly greater than *E*_short_.

## RESULTS

### Metabolic thermal response curves

The allometric scaling coefficient, α, and the apparent activation energy, *E*_*a*_, were consistent between the populations of *R. balthica* from the cold and warm streams (see Table 2 for model comparison and estimated parameter values). The temperature normalised rate of respiration, ln *b*(*T*_*c*_), and *T*_*h*_ (the temperature at which respiration was 50% inactivated) were both higher in the population of *R. balthica* from the warm stream. Because the optimum temperature, *T*_opt_, depends strongly on *T*_*h*_ (see Eq. (2)), *T*_opt_ was higher in *R. balthica* from the warmer stream (*T*_opt_ warm = 38.25 ± 0.6 °C; *T*_opt_ cold = 33.05 ± 1.5 °C). As ln *b*(*T*_*c*_) and *T*_opt_ were both higher, the warm populations of *R. balthica* had elevated metabolic rates across the full range of measurement temperatures (Fig. 2).

### Local adaptation of interaction strength

Interaction strength increased with elevated transplant temperature for the populations of *R. balthica* from both the warm and the cold streams (Fig. 3A; main effect of ‘transplant temperature’ GLM t_1,21_ = 2.56; *P* < 0.01). Furthermore, interaction strengths were consistently higher for the populations of *R. balthica* from the warm stream in both transplant temperatures (Fig. 3A; GLM main effect of ‘origin’ t_1,121_ = 2.92; *P* <0.005). These findings are consistent with the higher respiration rates observed in the warm population (Fig. 2) and highlight the association between metabolism and interaction strength.

### Disentangling the short- and long-term effects of warming on interaction strengths

Our experimental design enabled us to compare temperature sensitivities that capture short-term thermal acclimation (e.g. changes in interaction strength in response to the reciprocal transplant) as well as the long-term temperature sensitivity, which also includes effects of local adaptation (e.g. changes in rates between warm and cold populations quantified in the stream of origin). We found that interaction strength increased with temperature in both the short- and the long-term (Fig. 3). However, the magnitude of the temperature response was significantly larger in the long-term (Fig. 3; interaction between ‘transplant temperature’ and ‘timescale’ on interaction strength; GLM_1,18_ = -2.91; p < 0.05), where, the average *E*_short_ was 0.46 eV, while *E*_long_ was significantly higher at 0.99 eV. This divergence between the short- and long-term temperature sensitivities implies a non-trivial contribution of adaptation in amplifying the effects of temperature on interaction strength *in situ*, with the contribution of *E*_adapt_ of 0.51 and 0.53 eV in the cold and warm adapted populations respectively.

## DISCUSSION

Understanding how global warming will affect the strength of consumer-resource interactions and the stability of aquatic food webs is a fundamental challenge in evolutionary ecology that requires insight on the short-term effects of temperature on metabolism and interaction traits and they are modulated by evolutionary and developmental processes over longer time scales. There is evidence from terrestrial (Rall *et al.*, 2009; Barton, 2011; Vucic-Pestic *et al.*, 2011; Brose *et al.*, 2012), freshwater (Kratina *et al.*, 2012) and marine ecosystems (Sanford, 1999), that warming is likely to increase the strength of consumer-resource interactions, at least in the short-term, owing to the exponential effects of temperature on the consumption rates of mobile ectothermic consumers (Dell *et al.*, 2014; Gilbert *et al.*, 2014). What is less clear however, is how long-term responses to rising temperatures will modulate the direct effects of warming on species interactions. Space-for-time substitutions across broad spatial scales indicate that local adaptation to different thermal regimes can play an important role in shaping species interactions, often compensating for the direct effects of temperature on interaction traits (Barton, 2011; De Block *et al.*, 2012). Here, we build on this work by investigating the effects of temperature and local adaptation on the interaction between the gastropod grazer, *R. balthica*, and its algal resource. Our study contributes novel insights in a number of ways. First, we explore patterns of local adaptation over a relatively small spatial scale (m as opposed to km). The two streams in our experiment are separated by approximately 500 m but differ in temperature by 11 °C. Because dispersal, gene flow and genetic divergence among populations in this species are strongly related to geographic distance (Johansson *et al.*, 2016), our study over a relatively small spatial scale, provides insight into how metabolic and resource acquisition traits in closely related natural populations have divergeed in response to warming and is therefore directly relevant for understanding the effects of rapid climate change (Richter-Boix *et al.*, 2010; Keller *et al.*, 2013; Merilä & Hendry, 2014). Second, we quantified the effects of temperature on both metabolic and consumption rates to determine how temperature-driven selection on key traits shape the effects of long-term warming on the strength of consumer-resource interactions.

We found significant variation in the thermal response curves for respiration between the populations of *R. balthica* from the warm and cold streams. The optimum temperature (*T*_opt_) for respiration was higher in the warm population (i.e. metabolic rates peaked at higher temperatures). Furthermore, the inactivation energy (*E*_*h*_) was lower in the warm population, indicating that declines in the rate of respiration after the optimum (i.e. at high temperatures) were less pronounced than in grazers from the cold stream, where metabolic rates peaked at lower temperatures and declined markedly at temperatures above *T*_opt_. These divergences indicate that the different thermal regimes in these streams have selected for different metabolic traits in warm and cold populations of *R. balthica*. Whilst the higher *T*_opt_ and lower *E*_*h*_ in the warm population were in line with expectations assuming local thermal adaptation, we found no evidence that metabolic performance at high temperature was traded-off against performance at low temperature. Instead, metabolic rates were higher for *R. balthica* from the warm stream across all measurement temperatures. These results are in broad agreement with the “hotter is better” hypothesis, which proposes that maximal performance of organisms with higher optimal temperatures should be greater than those with lower optimum temperatures because of the thermodynamic constraints imposed by high temperatures on enzyme kinetics (Huey & Kingsolver, 1993; Kingsolver *et al.*, 2004; Angilletta *et al.*, 2010). Indeed maximal respiration rates in the population from the warm stream were greater than those from the cool (ln(R) warm stream: 3.39 ± 0.14 µmol O_2_ L^-1^ h^-1^, and cool stream: 2.54 ± 0.26 µmol O_2_ L^-1^ h^-1^, both ± 1.s.e.m.). The lower *E*_*h*_, (i.e. the steepness of the decline of the thermal reaction norm past the optimum), and higher ln *b*(*T*_*c*_), i.e. the rate of respiration normalised to 18 °C, in the warm population also meant that the thermal response curve for *R. balthica* from the warm stream was broader. In agreement with previous work (e.g. on bacteriophages, (Knies *et al.*, 2009), our data for the gastropod *R. balthica* indicate that adaptation to higher temperatures resulted in both greater maximal metabolic performance and a broader metabolic thermal reaction norm.

The general patterns observed in the metabolic traits were also reflected in the effects of temperature on interaction strength. Interaction strength was higher for individuals placed in the warm stream, irrespective of their stream of origin. These findings suggest that elevated temperatures increase consumption rates though the effects of temperature on respiratory physiology, but local adaptation to warmer environments also results in a correlated increase in metabolism and interaction strength at low temperature. This may have important wider implications for the effects of warming on the structure, functioning and stability of aquatic food webs (Rall *et al.*, 2009; O’Connor *et al.*, 2011; Vucic-Pestic *et al.*, 2011; Dell *et al.*, 2014; Fussmann *et al.*, 2014; Gilbert *et al.*, 2014). If long-term responses to increasing temperature give rise to higher maximal rates of metabolism and consumption as well as elevating rates at lower temperatures, then the effects of warming on the strength of consumer-resource interactions in the long-term could be greater than previously anticipated (Gilbert *et al.*, 2014). Indeed, work on experimental warming of aquatic ecosystems has shown that increases in the strength of top-down control can have profound effects on community structure and ecosystem processes (Burgmer & Hillebrand, 2011; Kratina *et al.*, 2012; Yvon-Durocher *et al.*, 2015). Elevated grazing rates at warmer temperatures can have a wide range of impacts in aquatic systems, with evidence for both increases (Yvon-Durocher *et al.*, 2015) and decreases (Burgmer & Hillebrand, 2011) in algal species richness, biomass and productivity.

In our experiments, the thermal sensitivities of metabolic rates were much larger than those of interaction strengths in the short-term (e.g. 0.96 and 0.45 eV respectively), in line with findings in other invertebrate systems (Rall *et al.*, 2009; Vucic-Pestic *et al.*, 2011; Fussmann *et al.*, 2014). These findings suggest that rates of grazing and metabolism were clearly linked, but became decoupled when individuals experience rapid changes in temperature that depart substantially from those in their local environment. In the short-term, if increases in metabolic demands with temperature are greater than those of consumption rates (as found here), then less energy will be transferred from the resource to the consumer,i.e. more is lost through respiration, see also (Rall *et al.*, 2009). If such imbalances are maintained over long periods of time then starvation of the consumer can ultimately result in a decline in top-down control on the resource (Fussmann *et al.*, 2014; Binzer *et al.*, 2015). However, when consumers’ feeding rates are more sensitive to temperature than metabolic rates, interaction strengths can become amplified in warmer environments, leading to faster resource depletion and eventually driving either the resource or the consumer to extinction (Vasseur & McCann, 2005). Long-term effects of temperature on interaction strengths have so far only been explored using food web models, parameterised using temperature sensitivities derived from short-term experiments (Vasseur & McCann, 2005; Rall *et al.*, 2012; Fussmann *et al.*, 2014). Consequently, such analyses don’t capture the evolutionary and developmental effects which can modulate the short-term effects of temperature on *per capita* rates. Our results highlight substantial differences between the short- and long-term effects of temperature on interaction strength; implying that longer term processes plays an important role in maintaining the balance between metabolic and consumption rates.

We quantified the short- and long-term effects of temperature in the reciprocal transplant experiment. The short-term temperature response (*E*_short_) captures the effects of physiological plasticity over the 48h experiment. Conversely, the long-term response (*E*_long_) also accounts for processes operating over longer timescales, including genetic micro evolution and non-genetic developmental effects of temperature. In our experiment, *E*_long_ was higher than *E*_short_, implying a significant role for long-term proceses in shaping the effects of temperature on *in situ* interaction strengths. Notably, the higher *E*_long_ was driven both by elevated grazing rates in the warm populations in the warm stream and lower rates in the cold populations in the cold stream. These results diverge from our expectations based on the metabolic cold adaptation hypothesis (Addo-Bediako *et al.*, 2000) which would predict populations from warmer environments should dampen the acute effects of temperature on metabolic rates. On the contrary, our results suggest that adaptation to warming amplified the effects of temperature on metabolic as well as grazing rates. The lower interaction strengths in the population of *R. balthica* from the colder stream highlight unexpected long-term effects of temperature on species interactions. Maintenance of lower than anticipated grazing rates in the cold stream could be selected for since lower grazing rates might result in greater food chain stability and/or stoichiometric homeostasis (Cross *et al.*, 2005; 2014) under the prevailing temperature regime. Thus, understanding the impacts of environmental change on the strength of consumer-resource interactions over timescales that are relevant to the rate of climate change (e.g. gradual warming over decades) will require an appreciation both of the direct effects of rising temperatures on species interactions and the reciprocal feedback between ecological and evolutionary dynamics (Fussmann *et al.*, 2007; Gravel *et al.*, 2010;Loeuille, 2010; Urban, 2013; Barraclough, 2015)

## Conclusions

We used a natural geothermal temperature gradient to investigate how warming influences the strength of algal-grazer interactions via the direct effects of temperature on metabolism and consumption, and indirect feedbacks through adaptation. Metabolic rates and interaction strength increased with temperature in the same way for both the warm and cold populations of *R. balthica*, suggesting that rapid changes in temperature have a consistent effect on interactions between mobile consumers and sessile resources, mediated by the effects of temperature on consumer metabolic rates (Dell *et al.*, 2014). However, the warm populations had higher metabolic and grazing rates across all measurement temperatures compared to their colder counterparts. These findings are consistent with the ‘hotter is better and broader’ hypothesis (Huey & Kingsolver, 1993; Knies *et al.*, 2009; Angilletta *et al.*, 2010) (e.g. adaptation to warming gives rise to higher maximal metabolic rates and broader thermal reaction norms). In consequence, our results suggest that warming could increase the strength of algal-grazer interactions, which are often ‘keystone’ interactions in aquatic systems, both via the thermodynamic effects of higher temperatures on enzyme kinetics and through correlated increases in *per capita* metabolism and consumption as organisms adapt to warmer temperatures.

## Conflict of interest

The authors declare no conflict of interest

## Acknowledgments

The authors thank Eoin O’Gorman and for comments on an earlier version of this manuscript. This study was funded by a Leverhulme Trust research grant (RPG-2013-335), and an ERC starting grant (ERC-StG 677278) awarded to GYD; and the University of Exeter.

## Supporting Information for Temperature-driven selection on metabolic traits increases the strength of an algal-grazer interaction in naturally warmed streams

**Figure S1:**
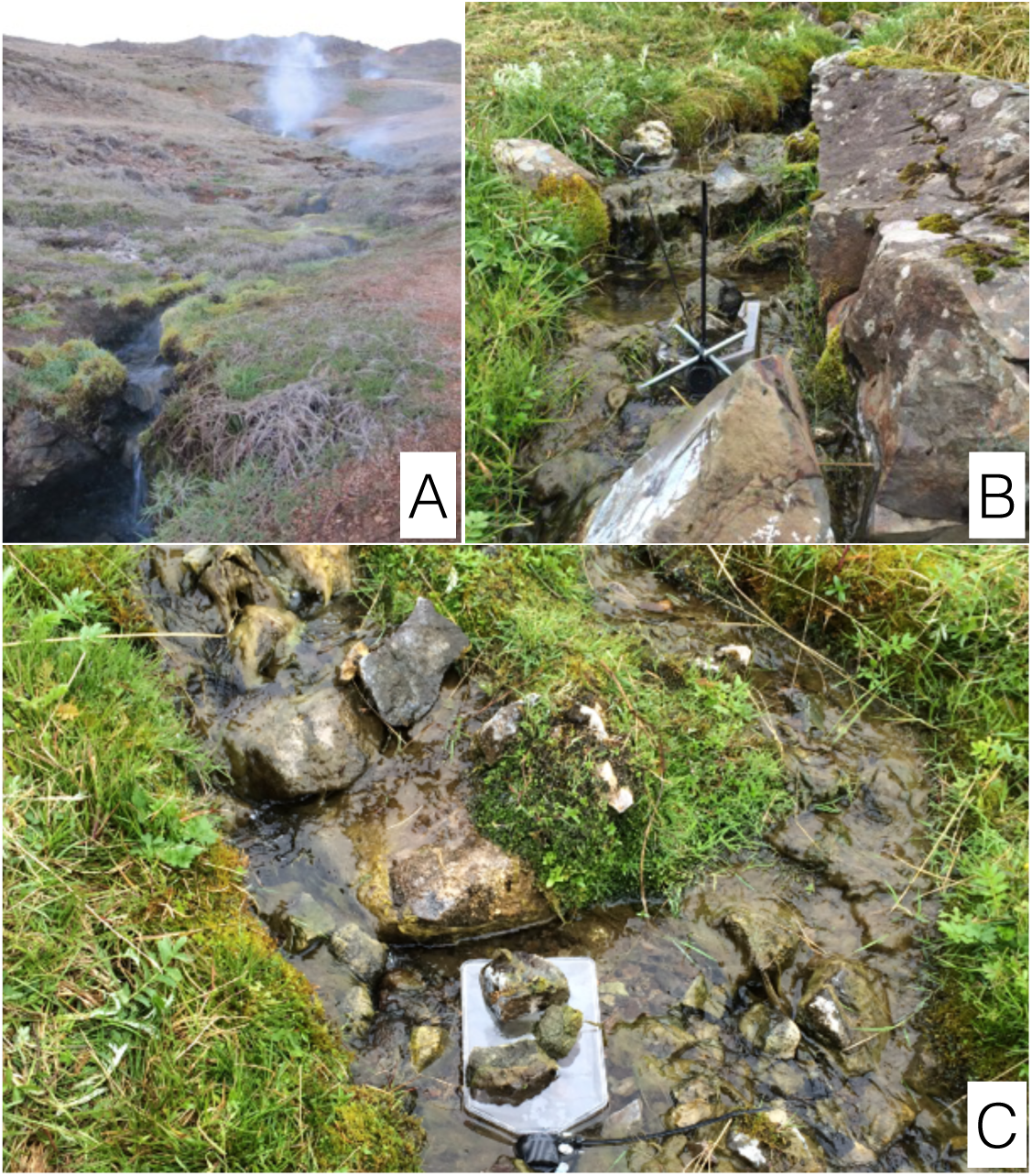
The catchment area and experimental set-up. A: View of representative stream in the Hveragerði catchment area. B and C: Microcosms fixed in stream 5 (Microcosm pictures courtesy of B. Flello).

**Figure S2:**
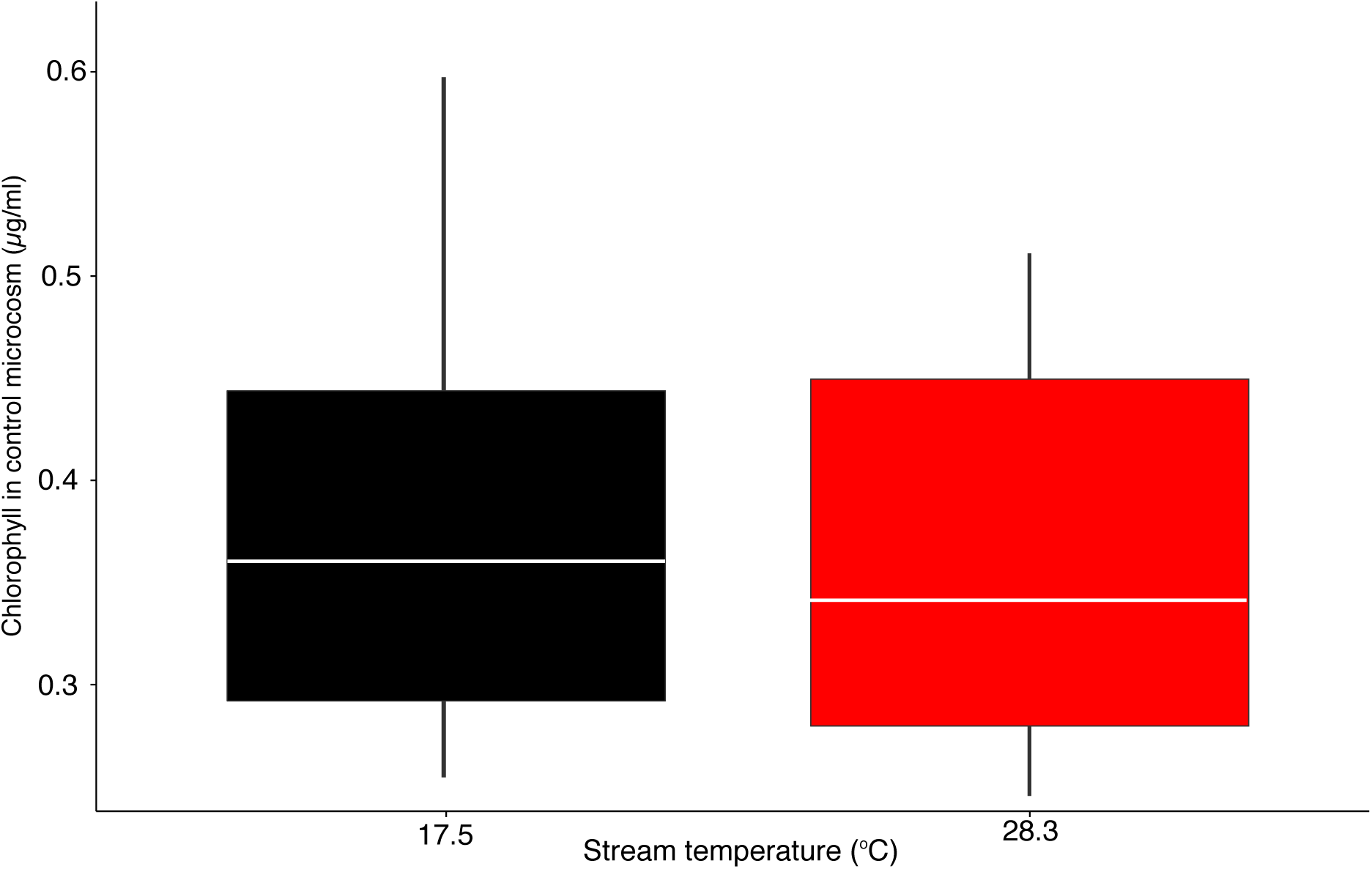
Chlorophyll content in control microcosms. Chlorophyll content at the end of the experiment in control microcosms did not differ significantly between cold and warm streams (one-way ANOVA F_1,10_=1.28, p=0.26) implying growth and mortality of the biofilms was consistent in both treatments.

**Figure S3:**
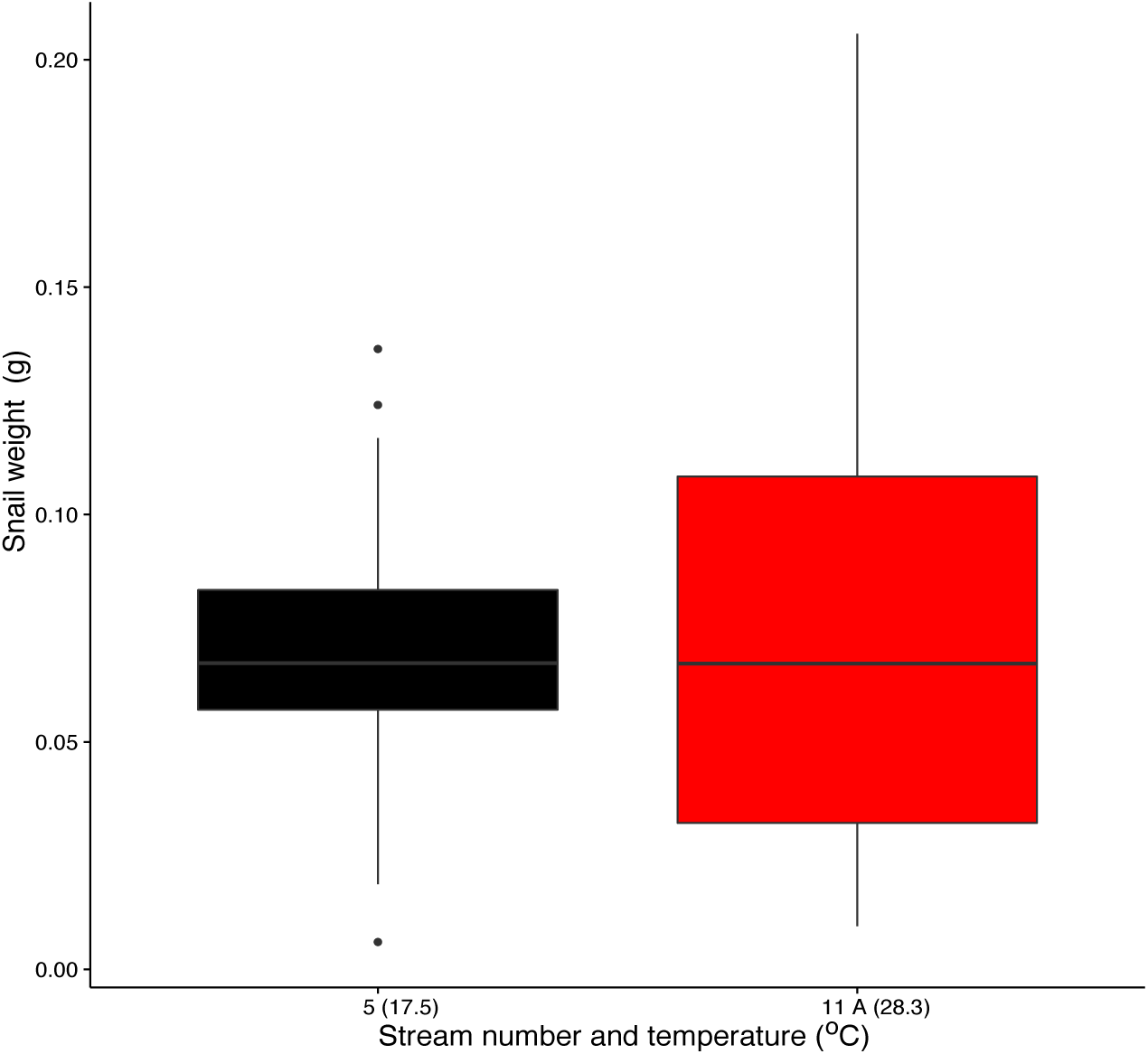
Snail weight for 205 randomly selected snails from each stream. Snail wet weight did not vary significantly between the cold (black) and warm (red) streams (one-way ANOVA: F_1,408_ = 0.15, p = 0.7).

**Figure S4:**
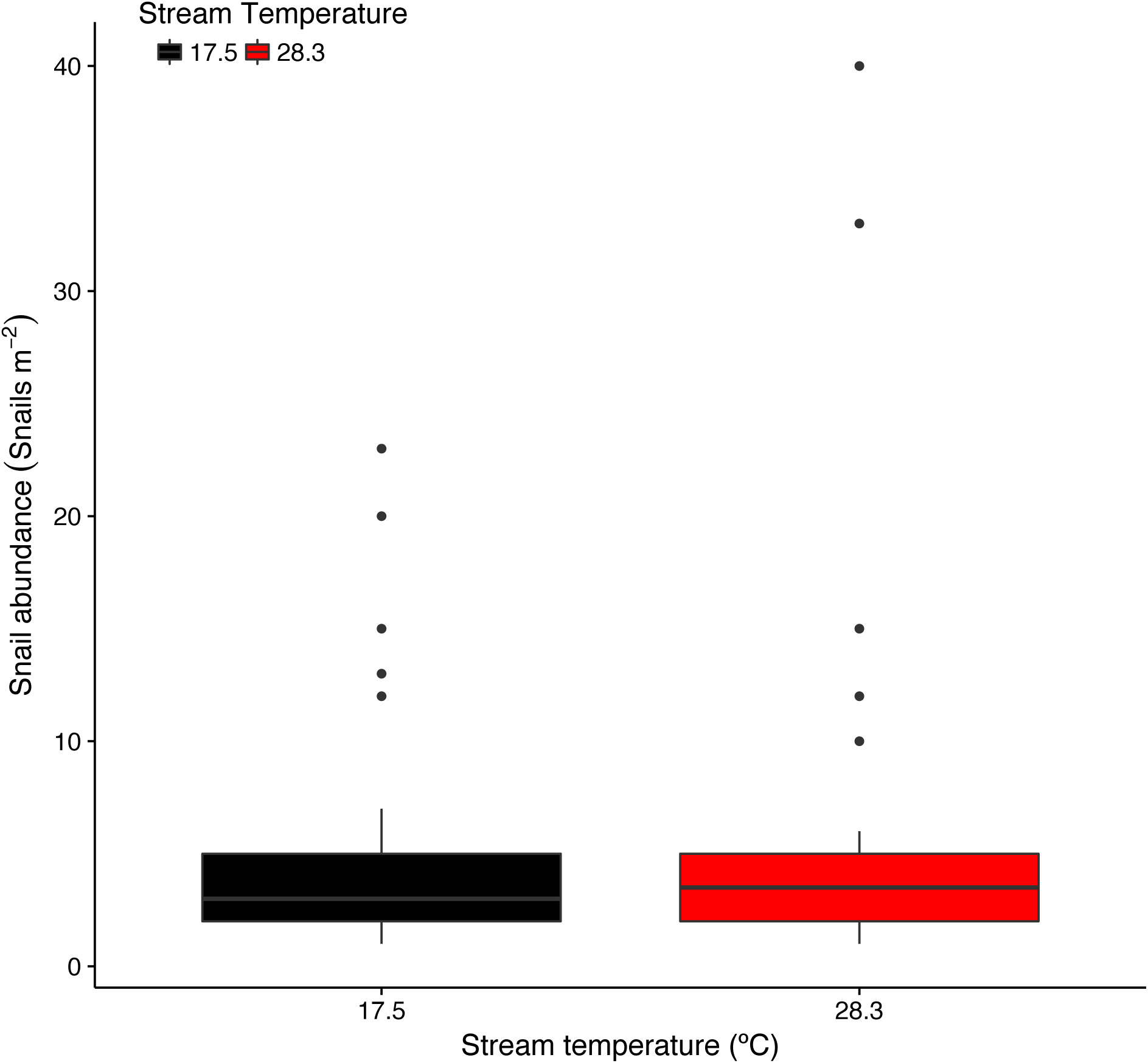
*In-situ* grazer density in the experimental streams. Snails were counted in 34 randomly distributed 1m^2^ quadrats. There was no significant difference in snail abundance between streams (one-way ANOVA: F_1,66_, p = 0.54). The mean abundance of 5.5 individuals per square meter was comparable to the densities established in the experimetal microcosms. Snails used for the experiments were sampled randomly from these patchess

**Table S1.**
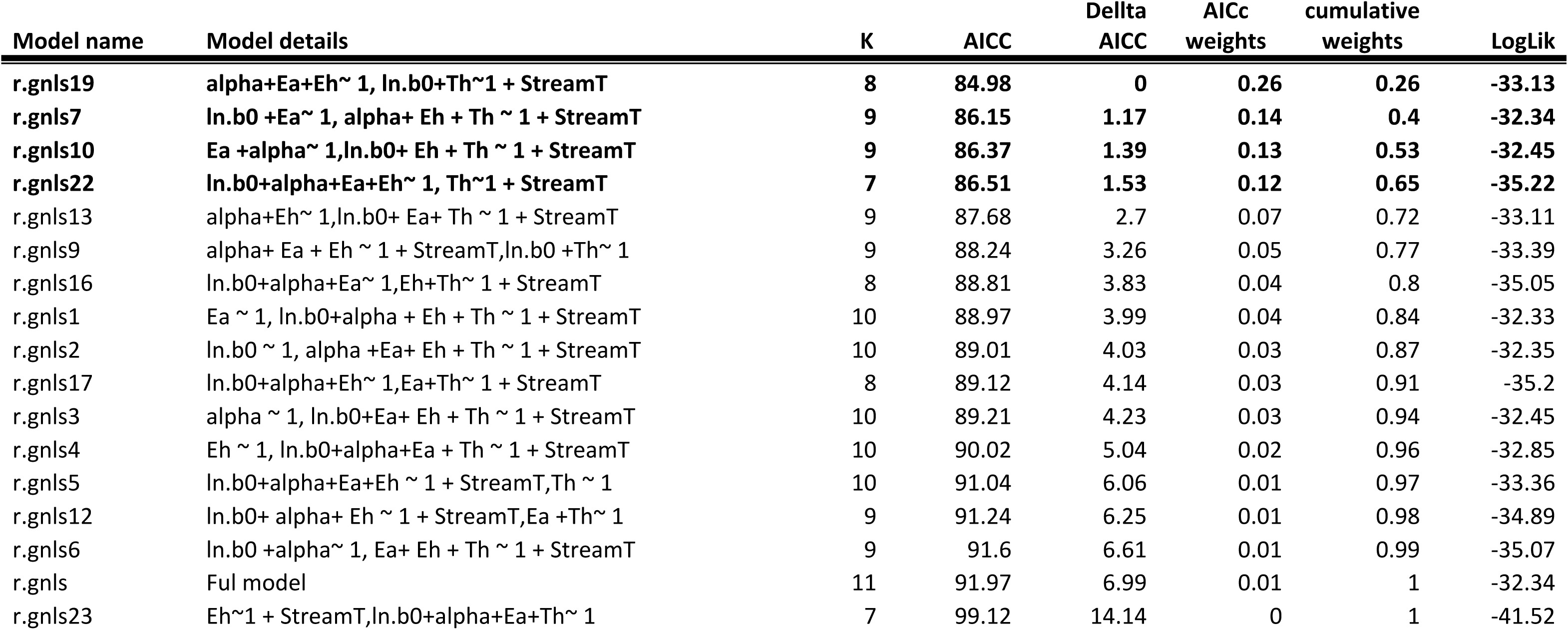
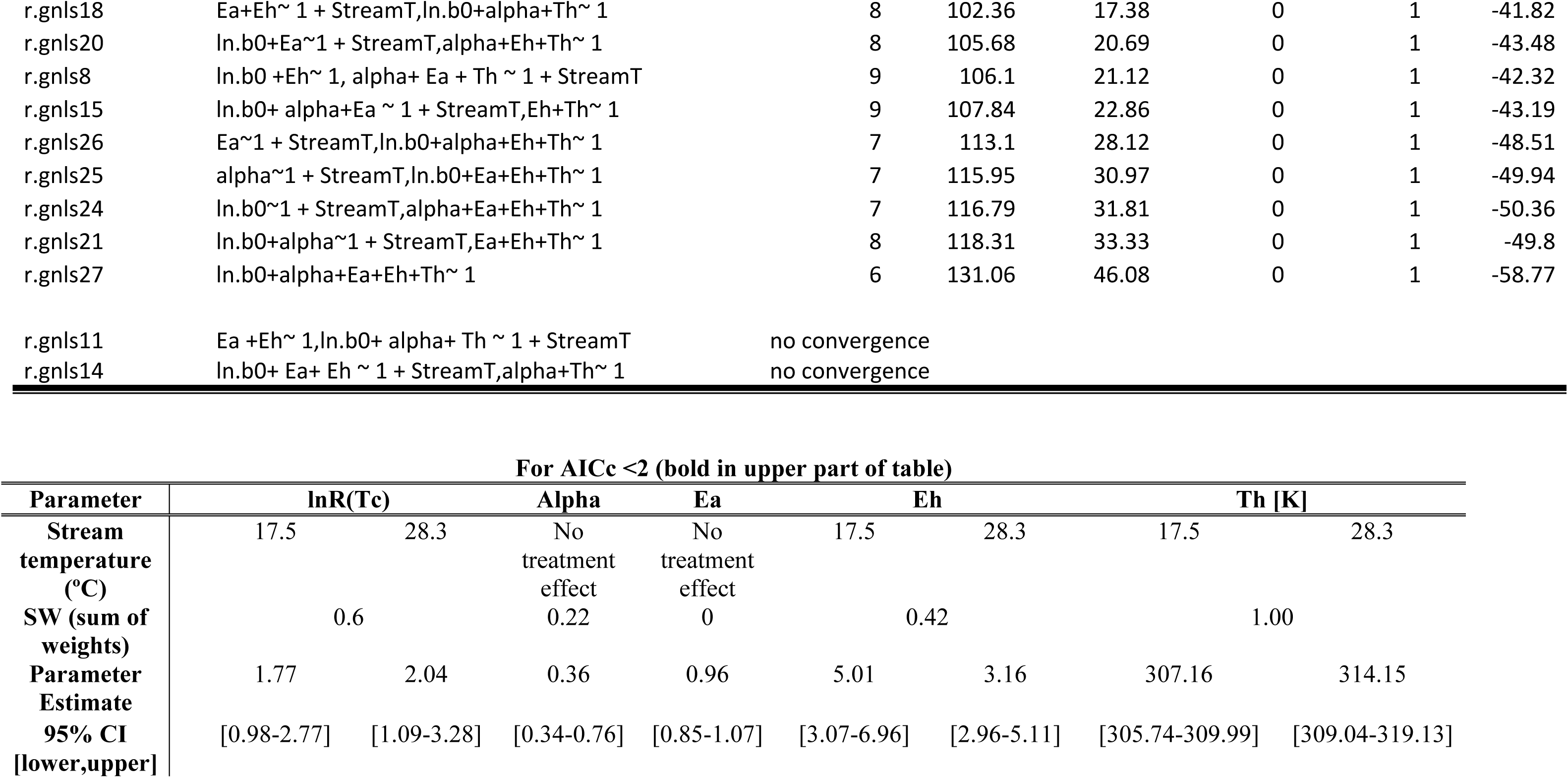
AICc comparison details for model simplification of the gnls models fitted to the thermal response data for respiration rates. Equation 1 was used to account for effects of mass (alpha in the model) on metabolic rate, and fitted to data for respiration (ln *R*) to characterize acute thermal responses. Model selection was carried out by fist fitting the most complex model to the data and then calculating AICc scores for all possible models with all possible treatment effects, and comparing the models through the aictab function within the AICmodav package (version 2.1-0) in R. The most parsimonious model included differences between warm and cold populations in ln *R*(*T*_*c*_), *T*_*h*_, and *E*_*h*_, while *E*_*a*_ and α were consistent between populations. For the four models with a difference in AICc of less than two, models were averaged using the modelavg function in the same package. **Lower part of table: Model averaged parameter estimates, with 95% confidence intervals (95%CI) and relative importance as sum of AICc based relative weights (SW), for models where Δ AICc <2**.

